# Sphingosine-1-phosphate cross-talks to Notch via a S1PR1-Dll4-MPDZ complex to regulate endothelial barrier function

**DOI:** 10.64898/2026.05.20.726610

**Authors:** Jennifer L. Bays, Jessica L. Teo, Freddy Suarez Rodriguez, Alanna M. Farrell, Amy E. Stoddard, Esther Koh, Timothy Hla, Christopher S. Chen

## Abstract

Sphingosine-1-phosphate (S1P) – a key bioactive component of high-density lipoproteins (HDL) – is instrumental in mediating HDL’s cardiovascular benefits, largely by enhancing endothelial barrier integrity^1, 2^. Here, we discovered that S1P induces Notch1 activation, and this Notch activation is required to enhance Rac1 activity and adherens junction assembly, which in turn stimulates endothelial barrier integrity. S1P rapidly activates Notch1 by stimulating the G-coupled protein receptor, S1P Receptor 1 (S1PR1) to drive internalization of the Notch ligand Delta-like protein 4 (Dll4). Notably, this internalization of Dll4 and subsequent activation of Notch does not involve traditional G-protein signaling; instead, S1P-bound S1PR1 forms a complex with Dll4 via the scaffolding protein MPDZ, and the undergoes co-endocytosis. Importantly, the loss or inhibition of Notch, Dll4, S1PR1, or MPDZ results in barrier defects. These findings elucidate a novel S1PR1-Dll4-MPDZ-Notch1 signaling axis that coordinates S1P and Notch signaling to regulate of endothelial cell signaling and barrier function.

## Main

The vascular barrier that separates blood from tissue is actively regulated by a dynamic, semipermeable endothelium that functions to preserve blood volume while permitting selective exchange of cells and nutrients. Disruption of this barrier is a key hallmark—and often a driving force—behind the etiology of both chronic and acute cardiovascular and cerebrovascular diseases, including retinopathies, atherosclerosis, stroke, and sepsis ^3, 4^. Barrier function is regulated primarily by adherens junctions (AJs), which disassemble during injury and reassemble during recovery^5, 6^. AJs are mediated by vascular endothelial (VE)-cadherin receptors, which in turn are anchored to the actin cytoskeleton via numerous scaffolding proteins^5^. Although endothelial barrier has long been known to be regulated by changes in cell-cell junctions, the extent to which different regulators of junction assembly might coordinate these processes remains unclear.

We previously identified Notch1 as a key driver of AJ assembly and barrier function, acting through a complex with VE-cadherin, Rac1, and the Rac GEF Trio^7^. Formation of this complex requires Notch1 activation, which depends on interactions with membrane-bound ligands on adjacent cells, followed by sequential cleavage events that activate the receptor^8, 9^. While we found that flow-dependent shear stress triggered Notch activity to drive barrier function, it remains unclear whether Notch acts solely in response to shear or if it integrates additional signals to modulate the barrier.

Bioactive lipids such as sphingosine-1-phosphate (S1P), a major component in the atheroprotective high-density lipoprotein (HDL) complexes, also stimulates Rac1 activation, AJ assembly, and vascular integrity—mirroring effects of Notch1^1, 2, 7, 10–12^. Stepping back, the parallels between S1P and Notch signaling extend beyond barrier function: both pathways also regulate endothelial quiescence, tip-stalk cell dynamics during angiogenesis, and atheroprotection^11, 13–17^. Based on these observations, we set out to investigate whether S1P and Notch1 may not operate as discrete regulators of Rac and junctions as traditionally assumed but instead overlap more intimately than previously recognized. Here, we demonstrate that S1P activates Notch1 signaling through a novel complex involving S1P Receptor1 (S1PR1), the Notch ligand Dll4, and the scaffolding protein MPDZ, revealing a previously unrecognized mechanism by which two distinct pathways are intimately engaged to regulate endothelial cell function.

## Results

### S1P activates Notch to regulate vascular endothelial barrier

To investigate whether S1P modulates Notch signaling, we treated primary dermal microvascular endothelial cells (MVECs) with a physiologic concentration of S1P (100nM) and assessed Notch1 activation, by the appearance of an epitope on the Notch1 intracellular domain (ICD) exposed upon γ-secretase-mediated cleavage^10, 18^. Notch1 cleavage was evident after one minute of S1P administration and remained elevated for 2 hours (Supplementary Fig. 1a), aligning with the ∼30 min half-life of S1P reported in cell culture^19^. This Notch activation was abolished upon treatment with γ-secretase inhibitor, DAPT (*N*-[*N*-(3,5-difluorophenacetyl)-L-alanyl]-*S*-phenylglycine *t*-butylester) or upon genetic deletion of *NOTCH1* via CRISPR-Cas9 (Fig. 1a, Supplementary Fig. 1b). The S1P-induced activation of Notch1 and its abolishment by DAPT were further validated by a Notch1 transcriptional reporter and by expression of Notch1 target genes, *HES1* and *HEY1* (Fig. 1b-c, Supplemental Fig. 1c). Notably, S1P-induced Notch activation was conserved across multiple human endothelial cell types, including lung, lymphatic, and aortic endothelia (Supplemental Fig. 1d-f).

**Figure 1.**
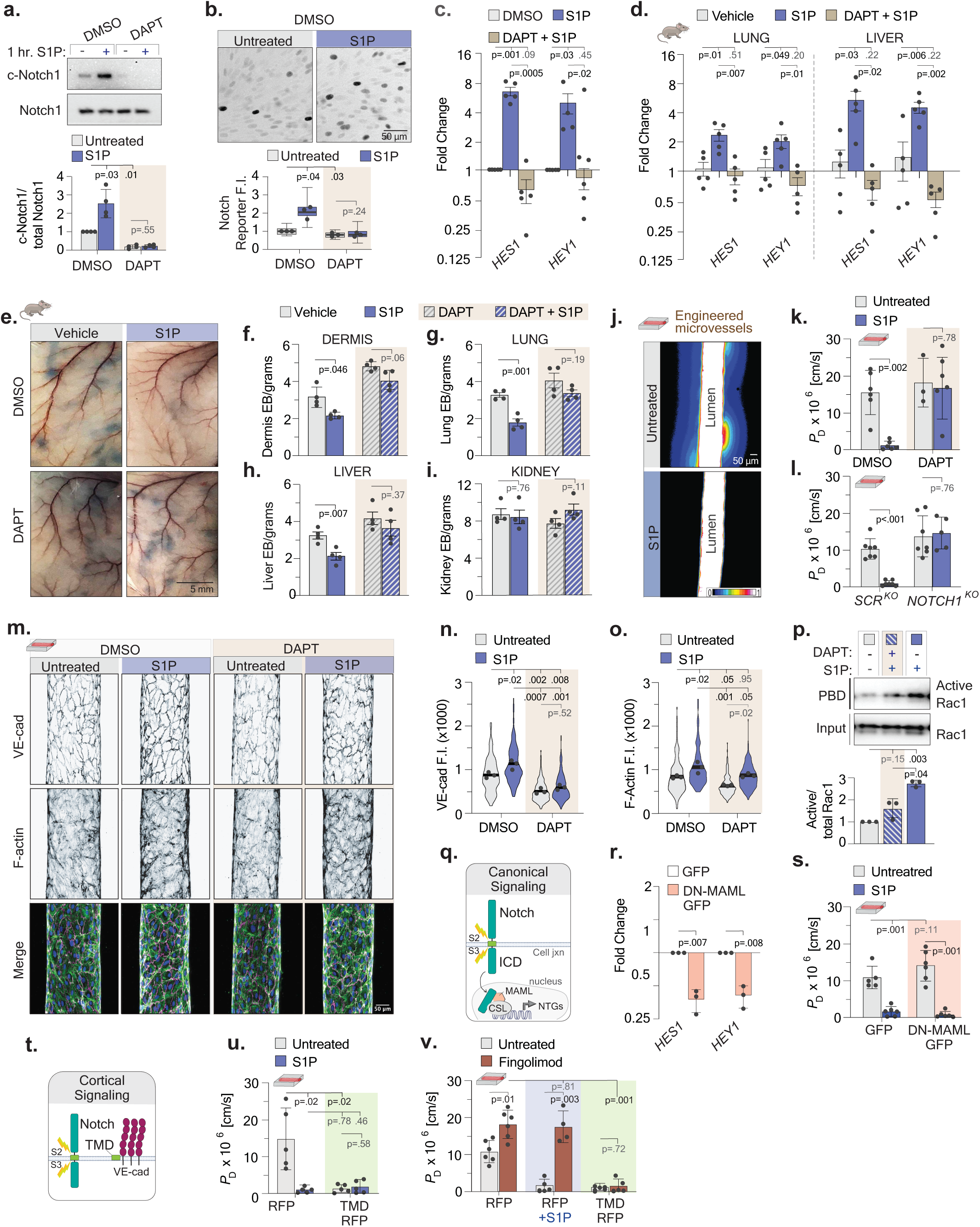
S1P activates Notch1 to enhance vascular endothelial barrier function. **a**, Immunoblots showing cleaved Notch1 (c-Notch1) and total Notch1 in MVECs treated with S1P for 1-hour ± γ-secretase inhibitor DAPT; quantification below (mean ± SEM). **b**, Fluorescent micrographs of YFP-Notch1 reporter intensity after S1P treatment; quantification below. Scale bar, 50 μm. **c-d**, Fold change in *HES1* and *HEY1* gene expression levels in (c) MVECs after S1P ± DAPT treatment, and in (d) mouse tissues after intravenous injection of S1P ± DAPT. **e**, Evans blue dye leakage in mouse dermis 1-hour post-injection with DMSO, S1P, or S1P + DAPT. Scale bar, 5 mm. **f-i**, Quantified dye leakage in (f) dermis, (g) lung, (h) liver, and (i) kidney, measured via absorbance. **j**, Heat maps of dextran dye diffusion in engineered microvessels ± S1P. Scale bar, 50 μm. **k-l**, Diffusive permeability in microvessels lined with (k) DAPT-vs. DMSO-treated or (l) *NOTCH1^KO^* vs. *SCR^KO^* MVECs (mean ± SD). **m**, Micrographs of VE-cadherin (magenta), actin (green), and nuclei (blue) staining in microvessels. Scale bar, 50 μm. **n-o**, Mean corrected intensities of junctional (n) VE-cadherin and (o) actin from (m). **p**, Rac1 activity (PBD pull-down). Active/total Rac1 quantification below. **q**, Schematic of canonical Notch signaling. **r**, *HES1*/*HEY1* gene expression levels in dominant negative-MAML-GFP (DN-MAML-GFP) normalized to GFP expressing cells. **s**, Permeability in DN-MAML-GFP vs GFP-lined microvessels ± S1P. **t**, Schematic of Notch cortical pathway mediated by its transmembrane domain (TMD). **u**, Permeability in TMD-RFP vs RFP-lined microvessels ± S1P. **v**, Permeability in TMD-RFP vs RFP-lined microvessels ± clinical S1P receptor degrader fingolimod.

We also confirmed S1P to activate Notch *in vivo*, as intravenous injection of S1P in mice significantly elevated *HES1* and *HEY1* expression in the descending aorta, lung, and liver—an effect that was abrogated by DAPT (Fig. 1d, Supplemental Fig. 1g-h). To examine if this S1P-induced Notch activity has a functional role, we examined S1P’s well-known regulation of vascular barrier integrity^10, 12^. We first confirmed vascular barrier effects with or without S1P, by examining co-injected Evans Blue dye extravasation in mice. As expected, S1P significantly reduced vascular leakage in the dermis, lung, and liver (Fig. 1e-h), Notably, co-administration of DAPT abrogated the protective effects of S1P (Fig. 1e-h), indicating that Notch cleavage is essential for S1P-mediated vascular barrier protection. Interestingly, we observed that S1P did not reduce vascular leakage in the kidneys (Fig. 1i), consistent with the absence of Notch induction in renal tissue in our studies (Supplemental Fig. 1h) and prior reports^20, 21^.

To begin to characterize this crosstalk mechanistically, we reproduced this S1P-Notch-barrier pathway in an engineered, endothelium-lined channel previously developed in our lab^7^. S1P significantly decreased transmural leak of fluorescence-tagged 70kD dextran in these engineered, with effects observed on timescales consistent with Notch1 cleavage (Fig. 1j, Supplemental Fig. 1a, i). To evaluate if Noth regulates barrier, we seeded engineered microvessels with CRISPR-Cas9-mediated Notch1 knockout cells or treated vessels with DAPT. Consistent with our *in vivo* data, inhibition of Notch abrogated S1P’s barrier-enhancing effects (Fig. 1k-l). In characterizing how S1P promotes barrier function in this model, we observed enhanced junctional VE-cadherin and cortical actin organization by quantitative immunofluorescence imaging, and Rac activity by pulldown assays (Fig. 1m-p), consistent with previous known effects of S1P on junctions and cytoskeletal signaling^10^. Interestingly, DAPT significantly inhibited the induction of Rac and junctional reinforcement by S1P (Fig. 1p). Together, these findings demonstrate Notch signaling to be a critical mediator of S1P-driven junctional remodeling and barrier function.

Given the S1P induces Notch cleavage and barrier within minutes (Supplemental Fig. 1a, i), we hypothesized that Notch regulates S1P-mediated barrier function through a non-canonical mechanism, rather than through canonical ICD-dependent transcriptional signaling. To test this, we seeded engineered microvessels with endothelial cells expressing a dominant-negative form of the transcriptional co-factor Mastermind-like protein 1^22^ (DN-MAML), which inhibits canonical Notch signaling. DN-MAML did not affect S1P-induced barrier enhancement, despite effectively suppressing Notch-dependent transcription (Fig. 1q-s). Similarly, overexpression of Notch ICD did not alter S1P barrier (Supplemental Fig. 1j-k).

We next investigated whether Notch enhances barrier function via a non-canonical, cortical signaling pathway. In this pathway, the transmembrane domain (TMD) remains embedded in the membrane following ICD shedding, where it binds to and reinforces VE-cadherin junctions and activates Rac signaling^7^ (Fig. 1t). Expression of the 33-amino-acid TMD alone was sufficient to enhance barrier integrity, recapitulating the barrier-enhancing effects of S1P (Fig.1u). These findings support a model in which S1P-induced Notch cleavage liberates the TMD, which then engages VE-cadherin to strengthen endothelial barrier function.

S1P receptors have been the target of a variety of new medications, including fingolimod (Gilenya®, Novartis), a widely prescribed treatment for relapsing multiple sclerosis^23^. Despite its efficacy, fingolimod has been associated with significant vascular complications in approximately 5% of its patients, including retinal leakage leading to macular edema, pulmonary edema, and peripheral lesions, likely due to fingolimod-induced degradation of S1P receptors^24–26^. We investigated whether activating cortical Notch signaling could counteract the barrier-disruptive effects of fingolimod. Within our engineered microvessels, we observe that fingolimod indeed compromised barrier integrity, which was not rescued by S1P administration, consistent with fingolimod inhibitory effect of S1P receptor signaling (Fig.1v). Excitingly, we find that activating cortical Notch signaling via expression of active Notch TMD protected barrier function in fingolimod-treated cells (Fig.1v). Together suggesting that Notch TMD signaling is downstream of S1P signaling required for endothelial barrier and could be a useful therapeutic target in setting of barrier disruption.

### S1PR1 activates Notch1 via Dll4

To investigate the mechanisms by which S1P activates Notch, we first focused on identifying which of the G protein-coupled receptors for S1P (S1PRs) might be involved^10^. Amongst the selective pharmacologic inhibitors for the three S1PRs expressed in endothelial cells (S1PR1, S1PR2, and S1PR3), only the non-specific S1PR degrader fingolimod and the S1PR1 inhibitor W146 significantly blocked S1P-induced Notch1 activation and barrier enhancement (Fig. 2a, 2d, 1v). These findings were corroborated using two independent siRNAs targeting *EDG1*, the gene encoding S1PR1 (Fig. 2b). Moreover, pharmacological activation of S1PR1 with the selective agonist AUY954^27^ was sufficient to induce Notch activation and barrier enhancement (Fig. 2c-d). Collectively, these findings establish a direct link between S1PR1 signaling and Notch activation.

**Figure 2.**
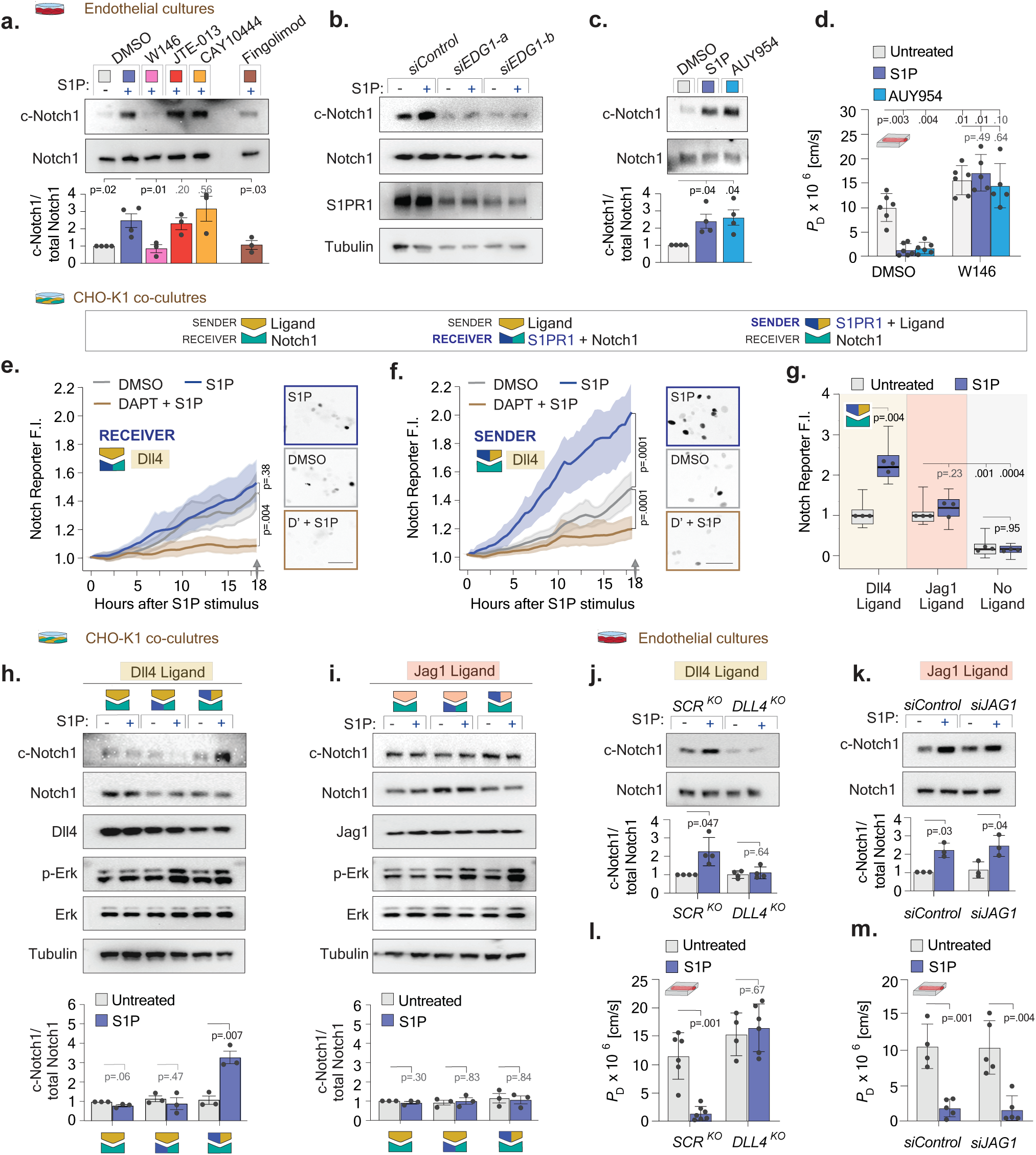
S1P stimulates S1PR1 in Dll4-, but not Jag1-, expressing sender cells to induce Notch signaling. **a**, Notch1 cleavage (c-Notch1) vs. total Notch1 in MVECs pretreated with inhibitors against S1PR1 (W146), S1PR2 (JTE-013), S1PR3 (CAY10444), or the non-selective S1PR modulator (fingolimod) before S1P stimulation. Quantification below (mean ± SEM). **b**, Immunoblots from MVECs treated with two separate siRNAs targeting *EDG1* (si*EDG1-*a, *siEDG1-*b) vs. non-targeting control (*siControl)*. **c**, Notch1 cleavage in MVECs treated with DMSO, S1P, or the S1PR1 agonist AUY954. Quantification below (mean ± SEM). **d**, Microvessel permeability ± S1P, AUY954, or S1PR1 inhibitor W146 (mean ± SD). **e-f**, Notch1 reporter activation in co-cultures of Notch1-expressing receiver cells and Dll4-expressing sender cells with S1PR1 in (e) receivers or (f) senders. Line graphs show reporter activity over time; micrographs and statistics at 18 h. Scale bar, 50 μm. **g**, Mean Notch reporter intensity in co-cultures with Dll4-, Jag1-, or no ligand-senders. Data normalized to untreated conditions per ligand; no-ligand data normalized to Jag1 untreated. **h-i**, Immunoblots from CHO co-cultures of Notch1-expressing receivers with (h) Dll4 or (i) Jag1 ± S1PR1 and S1P. Quantification below (mean ± SEM). **j-k**, Immunoblots of (j) MVECs with *DLL4* knockout (*DLL4^KO^*) or scramble control (*SCR^KO^*) and (k) *siJAG1* or *siControl*, ± S1P. Quantification below (mean ± SEM). **l-m**, Permeability in (l) *DLL4^KO^* vs. *SCR^KO^*-lined and (m) *siJAG1* vs *siControl*-lined microvessels, ± S1P.

Notch1 activation is classically mediated by endocytosis of Notch ligand in neighboring sender cells, triggering cleavage of ligand-bound Notch receptor in neighboring receiver cells^8, 9^. Given multiple Notch receptors and ligands are co-expressed in endothelial cells, each of which could potentially be modulated by S1PR1, we developed a reductionist system using Chinese hamster ovary (CHO) cells, which lack endogenous Notch, Notch ligands, and S1P receptor expression^28, 29^ (Supplemental Fig. 2a), permitting us to precisely dissect how S1PR1 may interface with the Notch pathway.

To identify the site of S1P action, we generated CHO lines to function as receiver or sender cells by expressing either the Notch1 receptor or Dll4, the prominently expressed Notch ligand in vasculature^30^. Expression of Notch or Dll4 alone was insufficient to confer S1P-induced Notch cleavage or activation of Erk, a canonical S1PR effector^12^ (Supplemental Fig. 2a). To render these cells responsive to S1P, we introduced a doxycycline-inducible, fluorescently tagged S1PR1 construct, titrated doxycycline to match expression levels in MVECs, and confirmed receptor activity by S1P-induced Erk phosphorylation (Supplemental 2b). Next, senders and receivers with or without S1PR1 were co-cultured at a 1:1 ratio and monitored for Notch transcriptional activity using a YFP reporter following exposure to S1P. Time-lapse imaging of co-cultures in which S1PR1 was expressed in Notch1-expressing receiver cells showed no difference in Notch activity between S1P-treated and vehicle-treated conditions (Fig. 2E and Supplemental Fig. 2c). However, when S1PR1 was expressed in Dll4-expressing sender cells, S1P stimulation increased YFP expression in the co-cultures (Fig. 2f and Supplemental Fig. 2d). DAPT reduced YFP expression in all conditions (Fig. 2e-f).

We then examined if S1P could induce Notch activity when employing another prominent endothelial Notch ligand, Jag1^8^. Surprisingly, Jag1, did not support S1P-induced Notch activation (Fig. 2g). Notch activation by S1P occurred only when S1PR1 and Dll4 were co-expressed in sender cells, a finding confirmed biochemically by detection of Notch cleavage (Fig. 2h-i). Importantly, S1P activated S1PR1 in all configurations, as evidenced by S1P-induced Erk phosphorylation (Fig. 2h-i).

To validate these findings in endothelial cells, we knocked down either Dll4 or Jag1. Loss of Dll4 expression significantly reduced S1P-induced Notch1 activation and barrier enhancement, whereas Jag1 knockdown had no effect (Fig. 2j-m, Supplemental Fig. 2e-f).

### S1PR1 triggers Dll4 endocytosis to facilitate Notch activation

Notch activation requires ligand endocytosis into sender cells to generate the mechanical pulling forces necessary for proteolytic receptor cleavage and activation^9^. Since S1P acts within sender cells, we investigate whether it influences Dll4 endocytosis. To monitor this, we employed a Dll4 construct tagged with CLIP, an enzyme-based labeling system that allowed us to selectively tag surface-exposed Dll4 just before S1P treatment^31^. S1P stimulation in endothelial cells significantly enhanced the number of Dll4 puncta, an effect that was abrogated by Dynasore, a dynamin inhibitor that disrupts vesicle scission (Fig. 3a-b, Supplemental Fig. 3a-b). Inhibiting dynamin also blocked S1P-induced Notch activation (Supplemental Fig. 3c-d), confirming the requirement of ligand endocytosis for Notch signaling. Next, we examined if S1PR1 specifically mediated Dll4 turnover. Knock-down of S1PR1 (encoded by *EDG1*) similarly impaired Dll4 endocytosis, highlighting S1PR1 as the principal receptor mediating this effect (Fig. 3a-b).

**Figure 3.**
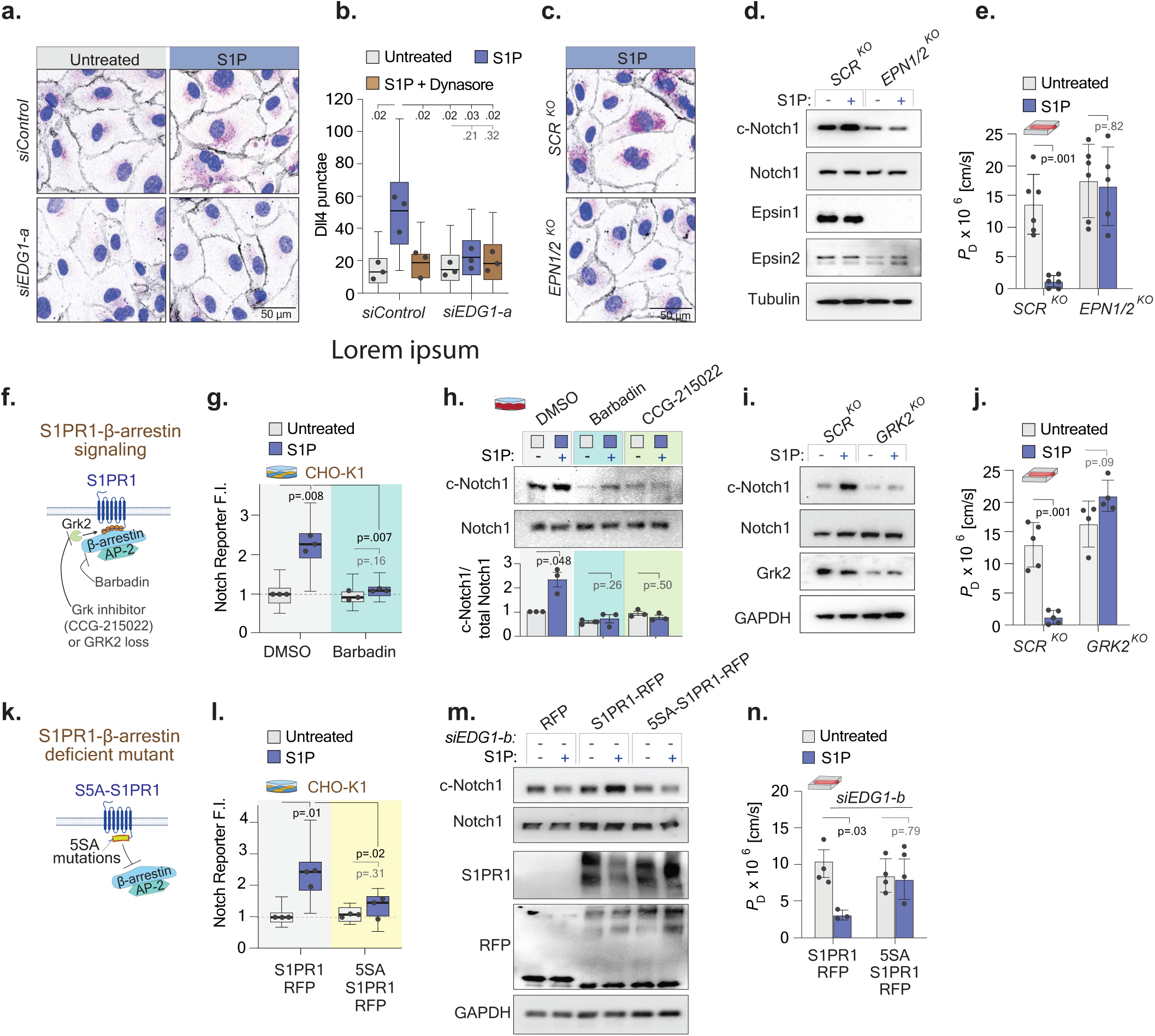
S1PR1 promotes Dll4 turnover and Notch activation via β-arrestin binding. **a**, Fluorescence micrographs of MVECs expressing CLIP-tagged Dll4, treated with *siEDG1*-a or *siControl*, labeled with cell-impermeable dye to mark surface Dll4 (magenta), and stained for β-catenin (gray) and nuclei (Hoechst, blue). Scale bars, 50 μm. **b**, Quantification of internal Dll4 puncta from (a); Dots represent mean values from independent experiments. **c**, Fluorescence micrographs of Dll4 labeling in MVECs expressing CRISPR-Cas9 guides for Epsin1/2 (*EPN1/2^KO^*) or scrambled control (*SCR^KO^*), labelled and stained as in (a). **d**, Immunoblots showing cleaved Notch and Epsin1/2 expression in *SCR^KO^* and *EPN1/2^KO^* MVECs. **e**, Permeability in *SCR^KO^* or *EPN1/2^KO^*-lined microvessels, ± S1P. **f**, Schematic of S1PR1 endocytic regulators. **g**, Notch1 reporter intensity after Barbadin or DMSO ± S1P. **h**, Cleaved Notch1 in MVECs pre-treated with Barbadin or the Grk inhibitor CCG-215022 ± S1P. **i**, Immunoblots of *GRK2^KO^* versus *SCR^KO^* MVECs, ± S1P. **j**, Permeability in *GRK2^KO^* or *SCR^KO^*-lined microvessels, ± S1P. **k**, Schematic of 5SA mutations disrupting β-arrestin binding to S1PR1. **l**, Notch1 reporter activity in CHOs expressing WT or 5SA-S1PR1 ± S1P. **m**, Immunoblots of MVECs with endogenous S1PR1 knockdown (*siEDG1*-b), rescued with WT S1PR1 or 5SA-S1PR1 ± S1P. **n**, Permeability in S1PR1 vs 5SA-S1PR1-lined microvessels, ± S1P.

Given that dynamin broadly regulates receptor turnover, we designed a genetic approach to target Dll4 endocytosis more specifically. We used CRISPR-Cas9-mediated deletion of *EPN1* and *EPN2*, which encode Epsin1 and Epsin2—endocytic adaptors that link Dll4 to the clathrin and AP-2 machinery^9^. Deletion of *EPN1*/*EPN2* blocked S1P-induced Dll4 internalization, Notch cleavage, and downstream endothelial barrier enhancement (Fig. 3c-e, Supplemental Fig. 3e-g). Together, these results demonstrate that S1P-S1PR1 signaling promotes Dll4 endocytosis in sender cells, which is essential for subsequent Notch activation.

We then examined whether S1PR1 turnover is similarly required for this process. S1PR1 internalization is triggered by G protein-coupled receptor kinase 2 (Grk2), which phosphorylates serine residues in the receptor’s C-terminus, recruiting β-arrestins that mediate endocytosis via the AP-2 complex^24, 32, 33^ (Fig. 3f). Treatment with Barbadin, a β-arrestin inhibitor that prevents AP-2 binding, blocked S1P-induced Notch activation (Fig. 3g)^34^. Consistent with this, both CRISPR-Cas9-mediated deletion of *GRK2* and pharmacological inhibition of Grk2 using CCG-215022 eliminated S1P-induced Notch cleavage and endothelial barrier enhancement (Fig. 3h-j).

Grk2 regulates the turnover of multiple GPCRs, necessitating a more targeted approach to assess S1PR1 endocytosis. To do so, we generated an endocytosis-deficient S1PR1 mutant (5SA-S1PR1), in which five Grk2-targeted serine residues were mutated to alanine^24, 33^ (Fig. 3k). Endothelial cells and CHOs alike expressing wild-type S1PR1 exhibited robust S1P-induced Notch activation, whereas those expressing 5SA-S1PR1 failed to activate Notch (Fig. 3l-m). Furthermore, 5SA-S1PR1 expressing endothelial cells were unable to enhance barrier integrity in response to S1P stimulus (Fig. 3n). These findings establish that endocytosis of both Dll4 and S1PR1 are required to link S1P signaling to Notch activation and vascular barrier regulation.

Many known functions of S1PR1 are mediated through canonical G-dependent signaling^29^. To assess whether this pathway is also required for S1P-induced Notch1 activation, we treated endothelial cells with pertussis toxin, which inhibits dissociation and downstream signaling of Gαi, the G-protein that couples with S1PR1 ^35^. While pertussis toxin effectively suppressed S1P-induced Erk phosphorylation, it had no effect on Notch1 cleavage or Notch reporter activity (Supplemental Fig. 3h-i). These results demonstrate that Gαi signaling is dispensable for S1P-induced Notch1 activation, supporting a distinct mechanism involving S1PR1 turnover rather than classical G protein-coupled signaling.

### S1PR1 forms a molecular complex with Dll4 via scaffolding protein MPDZ to facilitate Notch activation

Given that endocytosis of both Dll4 and S1PR1 are required for linking S1P signaling to Notch activation, we examined whether their internalization was coupled. To track S1PR1 trafficking, we generated a dual-tagged S1PR1 construct with an extracellular SNAP tag and intracellular mApple tag and expressed it in endothelial cells. Surface S1PR1 labeling confirmed its internalization and co-localization with mApple-S1PR1 and vesicle marker α-adaptin (Supplemental Fig. 4a-b).

To assess co-internalization with Dll4, we co-expressed SNAP-S1PR1 and CLIP-Dll4, labeling them with orthogonal dyes^31^. S1P stimulation induced co-endocytosis into puncta (Fig. 4a), prompting further biochemical investigation of their physical association. Co-immunoprecipitation revealed low basal binding of S1PR1 to Dll4, which increased upon S1P stimulation (Fig. 4b). This complex demonstrates specificity in that S1PR1 did not bind the Notch ligand Jag1 and Dll4 ligand did not bind S1PR3.

**Figure 4.**
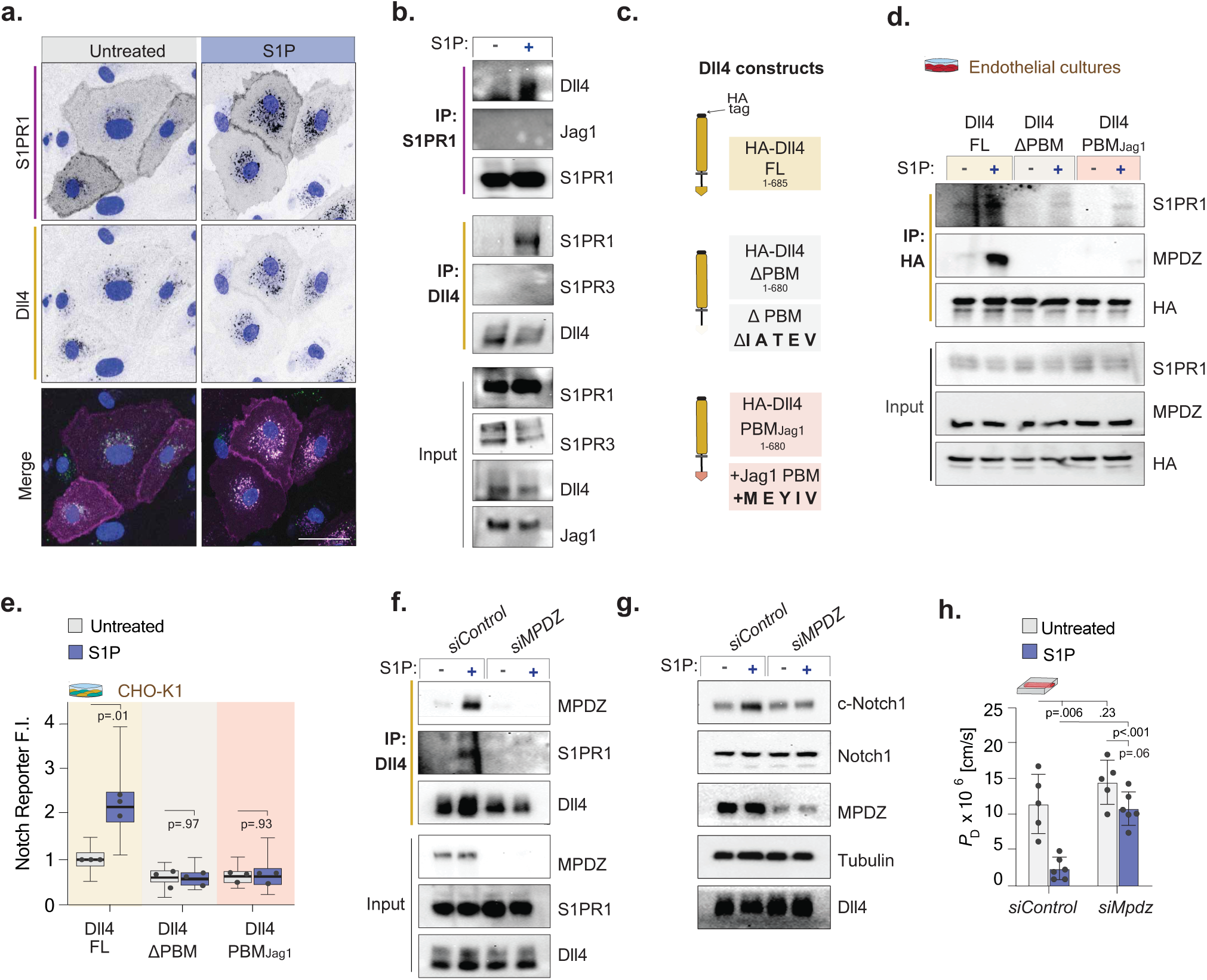
S1PR1 forms a complex with Dll4 and MPDZ to facilitate Notch activation and barrier function. **a**, Fluorescence micrographs of MVECs expressing SNAP-S1PR1 (magenta) and CLIP-Dll4 (green) ± S1P. Nuclei in blue. Scale bars, 50 μm. **b**, Co-Immunoprecipitation (co-IP) of Dll4 and S1PR1 from MVECs ± S1P, blots showing binding partners. **c**, Schematic of Dll4 constructs with PDZ Binding Motif removed (Dll4ΔPBM) or swapped with Jag1 PBM (Dll4-PBM_Jag1_). **d**, S1PR1 and MPDZ co-IP with HA-tagged Dll4 constructs ± S1P. **e**, Notch1 reporter intensity in CHOs expressing Dll4 constructs ± S1P. **f**, Co-IP of MPDZ and S1PR1 with Dll4 in MVECs treated with *siMPDZ* or *siControl*. **g**, Immunoblots of Notch1 cleavage and MPDZ expression in *siMPDZ* or *siControl* MVECs ± S1P. **h**, Permeability in *siMPDZ* or *siControl*-lined microvessels ± S1P.

To investigate the molecular basis and ligand specificity of this S1PR1-Dll4 complex, we focused on the cytoplasmic tails of these Notch ligands, which regulate endocytosis and protein interactions at cell-cell junctions^36^. Comparison of the intracellular domains (ICDs) of Dll4 and Jag1 revealed differences in their conserved PDZ-binding motifs (PBMs), located at the extreme C-termini. Dll4 contains a class I PBM consisting of the residues IATEV, whereas Jag1 harbors a class II PBM, MEYIV^36^. Deletion of the PBM in Dll4 (Dll4ΔPBM) or replacing Dll4’s PBM with that of Jag1 (Dll4-PBM_Jag1_) disrupted S1PR1-Dll4 complex formation as well as S1P-induced Notch1 activation (Fig. 4c-e, Supplemental Fig. 4c).

The PBM of Dll4 and Jag1 mediate specific interactions with unique PDZ-containing partners^36–38^. To identify the PDZ-containing partner that facilitates S1PR1-Dll4 interaction, we focused on MPDZ, a multi-PDZ domain scaffold implicated in endothelial Notch signaling^38^. Co-immunoprecipitation revealed that S1P stimulation enhanced MPDZ binding to both Dll4 PBM and S1PR1 (Fig. 4d, 4f). Given MPDZ’s established role in bridging membrane-associated complexes^39, 40^, we tested whether it links S1PR1 to Dll4. *MPDZ* depletion via siRNA reduced S1PR1-Dll4 binding and abolished S1P-induced Notch activation, without altering Dll4 expression (Fig. 4f-g). Finally, to assess the functional relevance of the S1PR1-MPDZ-Dll4 link, we examined its role in endothelial barrier regulation. *MPDZ* knockdown did not significantly alter basal permeability in engineered microvessels but abolished S1P-induced barrier enhancement (Fig. 4h). These findings establish MPDZ as a critical molecular link coupling S1PR1 to Dll4-mediated Notch signaling.

## Discussion

S1P and Notch1 have each been independently recognized as essential regulators vascular development and homeostasis. Yet, while S1PR1 and Notch1 share strikingly similar loss- and gain-of-function vascular phenotypes^1, 2, 7, 10–17, 41–45^, a direct link between them has been largely dismissed. Here, we identify S1P via S1PR1 is a critical regulator of Notch signaling by demonstrating that S1P-bound S1PR1 physically associates with Dll4 and promotes its endocytosis—revealing a previously unrecognized mechanism by which ligand internalization is initiated to drive Notch activation. Given the limited understanding of what drives Notch ligand endocytosis, our findings support a novel role for a GPCR as a trigger for Notch activation and may offer a broader framework for exploring ligand regulation in other contexts.

We further show that the Notch1 activity induced by S1PR1 is required for S1P to induce Rac activity, AJ assembly, and barrier function. Consistent with our prior work^7^, the cleavage of Notch receptor cleavage promoted barrier stability independent of its canonical transcriptional pathway, and instead relied on the release of the 33-amino acid Notch transmembrane domain (TMD) to locally activate Rac at cell-cell junctions. Notably, expression of the Notch TMD alone was sufficient to preserve barrier integrity in response to fingolimod-induced disruption^24–26^, highlighting its therapeutic potential in other vascular pathologies characterized by junctional insatiability.

While there is clearly substantial overlap between the S1P and Notch pathways, they likely contribute distinct functions as well given that loss of Notch transcriptional signaling fails to fully recapitulate the vascular defects associated with S1PR1 deletion^45^. Furthermore, S1PR1 deletion also has been reported not to significantly alter canonical Notch target gene expression in some settings^42, 45, 46^. Thus, while we have demonstrated a more intimate coupling between S1P and Notch signaling and its critical interdependence in barrier regulation, these pathways likely also serve distinct roles in certain developmental and adult contexts.

At the core of the interaction of these two pathways is the identification a novel S1PR1-MPDZ-Dll4 complex. While studies suggest a direction interaction between MPDZ and Dll4^36–38^, the precise mechanism linking MPDZ to S1PR1 remains unclear. Although S1PR1 lacks both a PDZ domain and a PDZ-binding site, it has been shown to bind PDZ-domain-containing proteins such as Prex-1, Trip-6, and Akt-1^47–49^, leading to the speculation that S1PR1 may contain internal, currently unmapped PDZ-binding motifs. Another possibility is MPDZ indirectly links Dll4 to S1PR1 through intermediaries such as β-arrestin, a mechanism observed with other PDZ-containing proteins^50, 51^. Regardless, it is plausible that other receptors and GPCRs may link with Notch ligands via PDZ domain-mediated interactions. If so, one would expect such complexes to be relatively selective given that the S1PR1-MPDZ-Dll4 complex could not accommodate either S1PR3 or Jagged1.

Lastly, it is noted that both S1P and Notch signaling also govern broader aspects of endothelial biology—including maintenance of quiescence, coordination of tip and stalk cells during angiogenic sprouting, and protection against inflammation and atherosclerosis^11, 13, 14, 17, 44^. Given these diverse roles, it would not be surprising if the regulatory mechanism described here may extend beyond vascular barrier function to influence additional critical vascular processes. Moreover, S1P receptor family members and Notch receptors are also co-expressed in other cells such as neurons and lymphocytes^52–54^, raising the possibility that coordinated signaling could also occur in even broader biological contexts. As such, the crosstalk between S1P and Notch described here underscores the need for deeper investigations into the possible functional interactions between these two important pathways.

## Figure legends

**Supplemental Figure 1. S1P induces Notch1 activation in multiple endothelial cells and tissues to regulate vascular barrier independent of Notch transcription.**

**a**, Notch1 cleavage in MVEC lysates following exposure to 100 nM S1P for the indicated time intervals (m, minutes; h, hours). Cleaved Notch1 (c-Notch1) and total Notch1 were assessed by western blot. The selected time point (1 hour) for subsequent assays is marked (&). **b**, Notch1 cleavage in lysates of *SCR^KO^*versus *NOTCH1^KO^* MVECs treated with or without S1P. **c**, Fluorescent micrographs of YFP Notch1 reporter expression (yellow) following S1P exposure, with or without DAPT; nuclei marked with H2B-BFP (blue). Scale bar, 50 µm. **d**, Fold change in *HES1* and *HEY1* gene expression levels in tissue-derived primary endothelial cells, from aortic blood endothelial cells (ECs), lung microvascular ECs and dermal lymphatics (mean ± SEM). **e-f**, Fold change in *HES1* and *HEY1* gene expression levels in (e) HUVECs and (f) the immortalized microvascular endothelial cell line HMEC-1. **g-h**, Fold change in *HES1* and *HEY1* gene expression levels in mice tissues from (g) descending aorta and (h) kidney tissues isolated from mice intravenously injected with S1P or co-treated with DAPT and S1P, normalized to vehicle-treated controls (n = 5, age- and sex-matched littermates). **i**, Diffusive permeability in engineered microvessels lined with wild-type (WT) MVECs treated with S1P for various time points or left untreated. The selected time point (1 hour) for subsequent assays is marked (&). **j**, Fold change in *HES1* and *HEY1* gene expression levels in cells expressing fluorescently tagged Notch1 ICD relative to fluorescent RFP control (mean ± SEM). **k**, Diffusive permeability in vessels lined with cells expressing ICD-RFP versus RFP control cells, with or without S1P treatment. RFP control vessels share values with those in Figure 1V.

**Supplemental Figure 2. S1PR1 expression enables S1P-induced Notch1 activation in a Dll4-dependent manner.**

**a**, Western blot showing expression of Notch1 receptor and ligands and that Notch cleavage and Erk phosphorylation are unchanged by S1P treatment in CHO cells expressing Notch1 or Dll4, either alone or in co-culture. **b**, Immunoblot showing titration of doxycycline to induce expression of S1PR1 in CHO cells to level comparable to endogenous S1PR1 in MVECs. The selected working dose (250 ng/µL) for subsequent assays is marked (&). **c-d**, Micrographs showing Notch reporter intensities in co-cultures of Notch1-expressing receiver cells and Dll4-expressing sender cells with S1PR1 in (c) receivers or (d) senders 18 hours after exposure to S1P, DMSO vehicle, or S1P + DAPT. Scale bar, 50 μm. **e**, Western blot verifying Dll4 knockout (*DLL4^KO^*) in MVECs using CRISPR-Cas9, compared to a scrambled control (*SCR^KO^*). **f**, Immunoblot confirming Jag1 knockdown in MVECs treated with Jag1-targeting siRNA (*siJAG1*) versus a non-targeting control (*siControl*).

**Supplemental Figure 3. Inhibition of dynamin and epsin impairs Dll4 internalization and suppresses Notch1 activation.**

**a**, Merged florescent images of MVECs expressing a dual-tagged (HA and CLIP) Dll4 construct, treated with *siEDG1-*a (S1PR1-targeting siRNA) or *siControl*, followed by treatment with the dynamin inhibitor Dynasore or DMSO vehicle ± S1P, as shown in Fig. 3a. Prior to S1P treatment, surface-exposed Dll4 was labeled with a cell-impermeable CLIP dye (magenta). After fixation and permeabilization, cells were stained with anti-β-catenin (gray) to mark cell boundaries and Hoechst (blue) to label nuclei. Scale bars, 50 μm. **b**, Individual fluorescent channels corresponding to the merged images in Figures 3A and S3A. After fixation and permeabilization, cells were stained with anti-HA (green) to visualize total Dll4, anti-β-catenin (gray) to mark cell borders, and Hoechst (blue) to label nuclei. Scale bars, 50 μm. **c**, Immunoblot of MVEC lysates treated with Dynasore or DMSO (vehicle) with or without S1P. **d**, Line graphs showing Notch reporter activation over time in CHO co-cultures treated with DMSO, S1P, S1P + DAPT, or S1P + Dynasore. Statistics done at 18 h mark. **e**, Merged Fluorescent micrographs corresponding to untreated conditions in Figure 3C. Scale bars, 50 μm. **f**, Florescent micrographs of *SCR^KO^*and *EPN1/2^KO^* expressing a dual-tagged (HA and CLIP) Dll4 construct, ± S1P. **g**, Quantification of internalized surface-labelled Dll4 in *SCR^KO^* and *EPN1/2^KO^* MVECs ± S1P. **h**, Immunoblot of MVEC lysates treated with pertussis toxin or vehicle, ± S1P. **i**, Box and whisker blot of Notch1 reporter fluorescence intensity (F.I.) following pertussis or vehicle treatment, ± S1P. Data normalized to vehicle/untreated controls.

**Supplemental Figure 4. Validation of S1PR1 and Dll4 PBM constructs.**

**a**, Fluorescence images of MVECs expressing SNAP-S1PR1-RFP, labeled with a cell-impermeable SNAP dye (green) before S1P treatment to mark surface S1PR1, compared to total S1PR1 (magenta). Scale bars, 20 μm. **b**, Fluorescence images of MVECs expressing SNAP-S1PR1, labeled for surface S1PR1 before S1P treatment, and immunostained for the endocytic protein α-adaptin. Enlarged regions (2× magnification) are shown in red boxes. Scale bar, 20 μm. **c,** Immunoblot showing lysates from CHOs expressing either hemagglutinin (HA)-tagged full-length Jag1 (Jag1 FL), or a modified Dll4 lacking its PDZ binding motif (Dll4ΔPBM) or in which its PBM was replaced with the PBM from Jagged1 (Dll4-PBM_Jag1_).

## Acknowledgements

We would like to thank J. Eyckmans and I.Z. Jaffe for scientific training, R. Honigsberg, A. Barutis, and J. Bardorf for technical assistance, and L. Kelleher for administrative support. For supporting the project, we thank NIH (HL147585, HL15507, HL155078, EB033821), The Paul G. Allen Frontiers Group Allen Distinguished Investigator Award, and the NSF Science and Technology Center for Engineering Mechanobiology. J.L.B acknowledges support from the NIH National Research Service Awards T32 EB16652 and F32 HL154664. J.L.T acknowledges support from American Heart Association Postdoctoral Fellowship 24POST1243689. F.S.R. acknowledges support from Research Council of Finland, decision number #337531 and #357911 and Instrumentarium Science Foundation. A.M.F. and E.K. from National Science Foundation Graduate Research Fellowship Program under Grant numbers 2234657 and 2141064, respectively. The opinions, findings, and conclusions or recommendations expressed are those of the authors and do not necessarily reflect the views of the funding agency.

## Author Contributions

J.L.B., J.L.T., A.M.F., and C.S.C. conceived and designed research; J.L.B., J.L.T.,F.S.R., A.M.F, and A.E.S. performed experiments; J.L.B., J.L.T., and E.K. analyzed data; J.L.B., T.H. and C.S.C. interpreted results of experiments; J.L.B. prepared figures; J.L.B. drafted manuscript; J.L.B., J.L.T., F.S.R., A.M.F, A.E.S, E.K., T.H. and C.S.C. edited and revised manuscript; J.L.B., J.L.T., F.S.R., A.M.F, A.E.S, E.K., T.H. and C.S.C.. approved final version of manuscript.

## Author Notes

During the preparation of this work the author J.M.B. used ChatGPT for grammatical editing. After using this service, the authors reviewed and edited the content as needed and takes full responsibility for the content of the publication. Correspondence: C. S. Chen.

## Methods

### Cell Culture

Human dermal microvascular endothelial cells (HMVECs), lung microvascular endothelial cells, aortic blood endothelial cells, and human umbilical vein endothelial cells (HUVECs) (all from Lonza) were cultured in EGM-2MV or EGM-2 growth medium (Lonza) and used between passages 2-8 (HDMECs, LMVECs, ABECs) or 2-10 (HUVECs). Immortalized microvascular endothelial cells (HMEC-1; ATCC) were maintained in EGM-2MV (Lonza). Cell line authentication (performance, differentiation, and STR profiling) was provided by Lonza.

Human embryonic kidney cells (HEK293T; ATCC) were cultured in Dulbecco’s Modified Eagle Medium (DMEM; Gibco) supplemented with 10% fetal bovine serum (FBS; R&D Systems) and 100 U/mL penicillin-streptomycin (Life Technologies). Chinese hamster ovary cells (CHO-K1; ATCC) were maintained in DMEM/F-12 medium supplemented with 10% FBS (R&D Systems) and 100 U/mL penicillin-streptomycin. CHO-K1 Notch1 reporter cells were a kind gift from Dr. John Ngo (Boston University). All cell lines were cultured at 37 °C in a humidified incubator with 5% CO₂. Mycoplasma contamination was routinely tested using the PlasmoTest mycoplasma detection kit (InvivoGen).

### Antibodies and Reagents

Antibodies for Western blotting (WB) and immunofluorescence (IF) were obtained from the following sources. From Cell Signaling Technology: cleaved Notch1 (Val1744, D3B8, 1:500 WB), total Notch1 (D1E11 XP, 1:1,000 WB), Dll4 (D7N3H, 1:1,000 WB), HA-tag (C29F4, 1:1,000 WB; 1:500 IF), Jagged1 (28H8, 1:1,000 WB), S1PR1 (E8U3O, 1:500 WB), Erk1/2 (137F5, 1:1,000 WB), phospho-Erk1/2 (Thr202/Tyr204, D13.14.4E, 1:1,000 WB), GRK2 (3982, 1:1,000 WB), GAPDH (D16H11 XP, 1:5,000 WB), β-Tubulin (2146, 1:5,000 WB), Vinculin (E1E9V XP, 1:5,000 WB), and HRP-conjugated secondaries (anti-mouse IgG, 7076; anti-rabbit IgG, 7074; both 1:5,000 WB). From Abcam: MPDZ/MUPP1 (EPR26317-59, 1:1,000 WB), RFP (ab124754, 1:1,000 WB). From Santa Cruz Biotechnology: S1PR1 (A-6, 1:100 WB), Epsin1 (C-11, 1:500 WB), Epsin2 (F-10, 1:300 WB), VE-cadherin (F-8, 1:300 IF), α-adaptin (C-8, 1:300 IF). From BD Transduction: Rac1 (610651, 1:200 WB), β-catenin (610153, 1:2,000 IF).

For immunofluorescence processing, Alexa Fluor 488-labeled phalloidin and Alexa Fluor Plus 488, 568, and 647 goat anti-mouse and anti-rabbit IgG highly cross-adsorbed (Life Technologies), were used at 1:100 and 1:1,000, respectively. DAPI was from Sigma-Aldrich. SNAP- and CLIP-tagged proteins were labeled with SNAP-Surface® 488 (S9124S, 5 μM) and CLIP-Surface® 647 (S9234S, 5 μM), purchased from New England Biolabs.

siRNA reagents were purchased from Horizon Discovery, including DharmaFECT 1 (T-2001), ON-TARGETplus Non-targeting Control Pool (*siControl*, D-001810-10-05), and ON-TARGETplus siRNAs for MPDZ (*siMPDZ*, J-019523-05-0005), JAG1 (*siJAG1*, L-011060-00-0005), and S1PR1 (si*EDG1-a*) vs. non-targeting control (*siControl* M-003655-02-0005). A custom ON-TARGET S1PR1 siRNA targeting the 3′UTR (si*EDG1-b,* CTM-523206) was also used with the following sequences: sense, 5′-AAGGAAAAGCUA-CACAAAAUU-3′; antisense, 5′-P-UUUUGUGUAGCUUUUCCUUUU-3′.

Small molecule reagents were from Cayman Chemical: sphingosine-1-phosphate (#62570, 100 nM), CCG-215022 (#31120, 1 μM), FTY720 phosphate (#10008639, 1 μM), doxycycline (#14422, 250 nM), JTE-013 (#10009458, 10 μM), CAY10444 (#10005033, 10 μM), and AUY945 (#9000548, 1 μM). From Sigma-Aldrich: DAPT (D5942, 20 μM), Barbadin (SML3127, 50 μM), Dynasore hydrate (D7693, 60 μM), W146 hydrate (W1020, 10 μM). Pertussis toxin (70323-44-3, 1 μg/mL) was from Millipore. All reagents were prepared and stored per manufacturer’s instructions.

### Mice experiments

All animal procedures complied with institutional guidelines and were approved by the Institutional Animal Care and Use Committee (IACUC). Male albino C57BL/6 mice (7-8 weeks old) were used for all in vivo studies.

To evaluate Notch activation, mice received a retro-orbital injection of 100 μL of 20 μM DAPT or vehicle (DMSO) in sterile PBS 16 hours prior to tissue harvest. The next day, mice were injected retro-orbitally with PBS containing S1P (0.03 mg/kg) along with 20 μM DAPT or vehicle. After 1 hour, mice were euthanized, and tissues were snap frozen.

For lung, liver, and kidney vascular permeability, the same injection protocol was followed with addition of Evans Blue dye (10 mg/mL). After 1 hour, blood was collected, mice were euthanized, and tissues were harvested, dried, and weighed. Dye was extracted using formamide.

To evaluate dermal vascular permeability, a Miles Assay was performed as previously described^1^. Briefly, mice were shaved 16 hours prior to the assay and received a retro-orbital injection of DAPT (20 μM) or DMSO. On the following day, pyrilamine maleate (8 μg/mL, 100 μL) was administered intraperitoneally to inhibit endogenous histamine release^2^. After 20 minutes, Evans Blue dye (10 mg/mL, 100 μL) containing S1P (0.03 mg/kg) with or without DAPT or vehicle was injected retro-orbitally. After 1 hour, blood was collected, mice were euthanized, and three flank skin sections were excised. Dye was extracted with formamide and quantified at 620 nm (reference 740 nm) using a Tecan Infinite M Plex plate reader. Results were normalized to blood dye content and tissue weight.

### Engineered microvessels

Three-dimensional engineered microvessels were fabricated using soft lithography and cast in polydimethylsiloxane (PDMS; Sylgard 184, Dow-Corning), as previously described^3^. Briefly, PDMS was mixed at a 10:1 ratio (base: curing agent), cured, and then cut and bonded to glass slides. Devices were treated with 0.01% poly-L-lysine followed by 1% glutaraldehyde, rinsed extensively with sterile water overnight, and sterilized with 70% ethanol for 30 minutes.

To form channels, sterile 160 μm-diameter steel acupuncture needles (Seirin) were inserted into each device. Devices were then UV-sterilized for 20 minutes. A 2.5 mg/mL collagen type I solution (prepared by diluting PureCol® Type I Collagen Solution, 3 mg/mL, bovine; Advanced BioMatrix) was buffered with 10× reconstitution buffer (250mM sodium bicarbonate, 200mM HEPES) and 10× DMEM (#D2429, Sigma-Aldrich) titrated to pH 7.5 before injection into the devices. Collagen was allowed to polymerize for 20 minutes at 37 °C. Growth medium was then added and incubated overnight. The following day, needles were carefully removed to create cylindrical microchannels (160 μm in diameter).

To seed endothelial cells, microvascular endothelial cells (MVECs) were detached using 0.05% Trypsin-EDTA, centrifuged at 200 × g for 5 minutes, and resuspended in EGM2-MV medium at 0.5 × 10⁶ cells/mL. Approximately 50-60 μL of the cell suspension was introduced into each device, and cells were allowed to adhere to the collagen matrix for 15 minutes before excess cells were washed out with fresh growth medium.

### Permeability Assays

Permeability within the engineered microvessels was assessed as previously described^3^. Briefly, 70 kDa Texas Red-conjugated dextran (Thermo Fisher Scientific) was introduced into the microvessels, and time-lapse imaging was performed using a Yokogawa CSU-21 spinning-disk confocal system mounted on a Zeiss Axiovert 200 M inverted microscope. Imaging was conducted with a Zeiss ×10 air objective and an iXon Ultra EMCCD camera (Andor). Images were acquired every 5 seconds for 5 minutes. Permeability coefficients were calculated from fluorescence intensity profiles using a published MATLAB script for image analysis^3^. Data are displayed as bar graphs of the average permeability ± standard error. Welch’s t-tests were used to compare vessel permeabilities between conditions.

### Immunofluorescence

For morphological analysis of adherens junctions and the actin cytoskeleton, cells were fixed and permeabilized with 1% paraformaldehyde and 0.03% Triton X-100 in PBS containing calcium and magnesium (PBS⁺) at 37°C for 90 seconds. Cells were immediately post-fixed in 4% paraformaldehyde in PBS⁺ at 37°C for 5–15 minutes. After fixation, cells were rinsed three times with PBS⁺ and further permeabilized with 0.1% Triton X-100 in PBS⁺ for 10 minutes. Blocking was performed with 2% BSA in PBS⁺. Primary and secondary antibodies were diluted in 2% BSA in PBS⁺. Cells were washed three times over 30 minutes with PBS⁺ between each antibody incubation step.

Immunofluorescent images were acquired using either a Yokogawa CSU-21 spinning disk confocal system mounted on a Zeiss Axiovert 200 M inverted microscope (Zeiss) equipped with a 40×/1.1 NA or 63×/1.15 NA oil-immersion objective and an iXon Ultra EMCCD camera (Andor), or using a Leica SP8 confocal microscope (Leica, Wetzlar, Germany) with a 10×/0.30 NA or 25×/0.95 NA water-immersion objective and Leica LAS X imaging software. Identical laser intensities and microscope settings were used across all samples within each experiment.

To quantify junctional VE-cadherin and actin intensities, a 12-μm line was drawn perpendicular to and centered on randomly selected cell-cell contacts. The Plot Profile tool in ImageJ was used to extract pixel intensity values along each line. Profiles were captured for 60 independent junctions obtained from three separate vessels. Images were corrected for background fluorescence. Two-way ANOVA used for Statical comparisons using Prism 10.

For endocytosis assays, endothelial cells (ECs) expressing CLIP-tagged Dll4 and/or SNAP-tagged S1PR1 were seeded onto glass slides and cultured for 48 hours. Cells were treated with either 60 µM Dynasore hydrate or DMSO (vehicle control) for 60 minutes prior to labeling. CLIP- and SNAP-tagged proteins were labeled using CLIP-Surface® 647 (5 μM, New England Biolabs) and/or SNAP-Surface® 488 (5 μM, New England Biolabs), respectively, in media containing Dynasore or DMSO for 10 minutes at 37°C. After labeling, cells were washed with Dynasore- or DMSO-containing medium and cultured statically for 45 minutes before fixation with 4% paraformaldehyde in PBS⁺ at 37°C for 15 minutes.

Cells were then permeabilized with 0.025% saponin and 2% BSA in PBS⁺ for 30 minutes at room temperature. All subsequent antibody incubations and washes were performed in this buffer. Cells were incubated overnight at 4°C with mouse anti-β-catenin (1:2000) and rabbit anti-HA (1:200) antibodies. The next day, cells were incubated with AlexaFluor-488-conjugated anti-mouse (1:200) and AlexaFluor-568-conjugated anti-rabbit (1:200) secondary antibodies for 2 hours at 4°C. Internalized Dll4 was quantified by applying a threshold and using the particle analysis tool in ImageJ. Mean reporter intensity per FOV was normalized to the untreated condition. The range of Dll4 puncta/cell is shown with box-and-whisker plots, and the average number of Dll4 puncta/cell from each experiment overlaid as individual dot plot. Welch’s t-tests were used to compare the average number of Dll4 puncta/cell from each experiment.

### Cloning

A Flag-tagged EDG1 cDNA (encoding the S1PR1 protein), containing an N-terminal cleavable signal sequence to promote membrane localization, was obtained from Addgene (plasmid #66496). This construct was modified to include a C-terminal mApple fluorescent tag and cloned using Gibson assembly into either a third-generation lentiviral pRRL CMV expression vector or the inducible pCW57-tetON lentiviral vectors (Addgene plasmids #71782 and #89180), which confer puromycin or neomycin resistance, respectively. A phosphorylation-deficient mutant, S5A-S1PR1, was generated by mutating five C-terminal serine residues to alanines (S351A, S353A, S355A, S358A, and S359A) using Gibson assembly. To enable surface labeling, we added a SNAP-tag which is derived from *O6-alkylguanine-DNA alkyltransferase* and reacts with benzylguanine (BG)-conjugated substrates. The SNAP-tag was a kind gift from Dr. John Ngo (Boston University) and was inserted on the N-terminus between the signal peptide and the Flag-tag in S1PR1 constructs.

Human DLL4 cDNA (Sino Biological) was modified by PCR to include an N-terminal hemagglutinin (HA) tag and cloned into a third-generation lentiviral pRRL CMV expression vector. For endocytosis assays we added a CLIP-tag, which is an engineered variant of O6-alkylguanine-DNA alkyltransferase that reacts with benzylcytosine (BC)-conjugated substrates. This CLIP-tag (Addgene #101136) was inserted into the DLL4 construct between the extracellular domain (after residue P524) and the transmembrane domain (before V530) of DLL4. For compatibility with CRISPR/Cas9 DLL4-targeted cells, the sequence within the construct targeted by our CRISPR guide target sequence was modified using Gibson Assembly cloning from ttC-ATC-AAC-GAG-CGC-GGC-GTA-Ctg to ttT-ATT-AAT-GAA-AGA-GGA-GTC-Ttg, preserving the encoded amino acid sequence as FINERGVL (aa 37-44). To generate a DLL4 construct lacking the PDZ-binding motif (DLL4ΔPBM), the final 15 base pairs of the DLL4 coding sequence were deleted. For the DLL4-swPBMJag1 chimera, the last 15 base pairs of DLL4 were replaced with the PDZ-binding motif sequence *MEYIV* from Jag1. Jag1 cDNA was amplified from Addgene plasmid #17336 and cloned into a modified third-generation lentiviral pRRL CMV expression vector. The reciprocal chimera, Dll4-PBM-Jag1, was created by replacing the final 15 base pairs of Jag1 with the DLL4 PDZ-binding motif sequence *IATEV*.

To generate a Notch transmembrane construct, the sequence spanning the S2 to S3 cleavage site (Val1721-Gly1753) was synthesized (IDT) with EcoRI and AgeI restriction sites and cloned into the multiple cloning site (MCS) of a third-generation lentiviral pRRL CMV expression vector containing a C-terminal mApple tag. To generate a Notch intracellular domain (NICD) construct, the region spanning the S2 cleavage site through the C-terminal end of the intracellular domain (Val1721-Lys2555) was PCR amplified and inserted upstream of the mApple tag in the same vector. These constructs were expressed via lentiviral transduction. The dominant-negative MAML1 construct (dnMAML; MSCV-IRES-GFP MAML1 13–74) was a kind gift from Dr. Jon Aster.

### Quantitative PCR

Endothelial cells were lysed with Trizol (LifeTechnologies) followed by chloroform phase separation. The aqueous phase was collected and diluted with 70% ethanol at a 1:1 ratio, loaded on an RNA micro column (RNA microkit, Qiagen Sciences) and RNA extraction was performed according to the manufacturer’s protocol. Subsequently, 0.8-1 μg total RNA was converted to cDNA with qScript cDNA Supermix (Quanta Sciences). Real-time PCR was performed in 20-μl reactions using the PowerUp SYBR Green Mastermix (AppliedBiosystems) and a QuantStudio3 (ThermoFischer), with 40 cycles of 95°C for 15 seconds and 60°C for 1 minute followed by a melt curve of 95°C for 15 seconds and 60°C for 1 minute and 95°C for 1 second. Relative gene expression is expressed as 2–ΔΔ*C*t, in which Δ*C*t is the difference in *C*t value between the gene of interest (GOI) and the housekeeping gene (proteasome subunit beta type-2 (*PSMB2*) and ΔΔ*C*_t_ is the difference between the Δ*C*_t_ of the GOI in an experimental condition and the Δ*C*t of the same GOI in a control condition. qPCR Primer sequences for human genes are *HES1* [Forward-CCAAGTGTGCTGGGGAAGTA, Reverse-CACCTCGGTATTAA-CGCCCT], *HEY1* [Forward-CTGAGCAAAGCGTTGACA, Reverse-TCCACCAACACTCC-AAA], *PSMB2* [Forward-ACTATGTTCTTGTCGCCTCCG, Reverse-CTGTACAGTGTCTC-CAGCCTC]. qPCR Primer sequences for mouse genes are *HES1* [Forward-CAACACGACAC-CGGACAAACCAAA, Reverse-TGGAATGCCGGGAGCTATCTTTCT], *HEY1* [Forward-CGCGGACGAGAATGGAAACT, Reverse-GCCAAAACCTGGGACGATGT], *PSMB2* [Forward-CGCTTCATCCTGAATATGCCA, Reverse-TTAGGAACCCTGTTTGGGGAAG], and GAPDH [Forward-AGGTCGGTGTGAACGGATTTG, Reverse-TGTAGACCATGTAGT-TGAGGTCA]

### Lentiviral-Mediated CRISPR Genome Editing

CRISPR knockout (KO) cells were generated using the lentiCRISPRv2 system (a gift from F. Zhang; Addgene plasmid #52961). Guide RNAs (gRNAs) were designed using the CRISPOR tool^4^ and cloned into the BsmBI site of the lentiCRISPRv2 plasmid. gRNA sequences are Scramble (GCACTACCAGAGCTAACTCA), NOTCH1 (CGTCAGCGTGAGCAGGTCGC), DLL4 (CATCAACGAGCGCGGCGTAC), EPN1 (GGCGGAGATCAAGGTTCGAG), EPN2 (CATCGTGAACAATTACTCAG), and GRK2 (TCAGTGGCACTCTTCGAGAA). Lentiviral particles were produced by co-transfecting HEK-293T cells with the gRNA-containing lentiCRISPRv2 plasmid and the packaging plasmids pVSVG, pRSV-REV, and pMDL using calcium phosphate transfection. Viral supernatants were collected 48 hours post-transfection, concentrated using PEG-it™ Viral Precipitation Solution (System Biosciences), and resuspended in PBS. Target cells were transduced overnight in growth medium and selected with 2 µg/mL puromycin starting 48 hours after infection. CRISPR-Cas9-mediated knockouts were validated by Western blot.

### siRNA Transfection

Endothelial cells were seeded into 6-well plates in antibiotic-free EGM2-MV medium 24 hours prior to siRNA transfection. Transfection reagents and siRNA were diluted in Opti-MEM (Thermo Fisher Scientific) and added to wells containing cells at approximately 80% confluency, following the manufacturer’s protocol. siRNAs were applied at a final concentration of 40 nM. After 24 hours, cells were switched to assay media and used for downstream applications. For quantification of knockdown efficiency, cells were lysed for western blot analysis 72 hours post-transfection.

### Cell Lysis and Western Blotting

Cells were lysed either directly in 2× sample buffer (0.2 M Tris-HCl, pH 6.8, 4% SDS, 20% glycerol, 0.03% bromophenol blue, and 5% β-mercaptoethanol) or in RIPA buffer supplemented with 2× protease and phosphatase inhibitors (Thermo Fisher Scientific, 78442). Lysates were homogenized by passing through a 25-gauge needle three times, then clarified by centrifugation at 15,000 g for 15 minutes. Protein concentration was determined using the Pierce BCA Protein Assay Kit (Thermo Fisher Scientific, 23225) and diluted in 2× sample buffer. Proteins were separated by SDS-PAGE using NuPAGE™ Bis-Tris or Tris-Acetate gels (Thermo Fisher Scientific), run at 180–220 V in MOPS running buffer, and transferred to PVDF membranes using NuPAGE transfer buffer at 250 mA for 90 minutes. Membranes were blocked in TBS-T containing 5% nonfat dry milk for 1 hour, followed by incubation with primary antibodies diluted in blocking buffer overnight at 4 °C on a rocker. After three washes at room temperature, membranes were incubated with HRP-conjugated secondary antibodies (1:5,000) for 1 hour, washed three times, developed with SuperSignal™ West Dura Extended Duration Substrate (Thermo Fisher Scientific), and imaged using the iBright™ CL1500 Imaging System (Thermo Fisher Scientific). Band intensities were quantified by integrated density using ImageJ, background-subtracted, normalized to a loading control for each sample. Samples were then normalized to the untreated conditions and statistical comparisons made using Welch’s t-tests.

### Rac activity assay

PAK-PBD beads to bind active Rac1 (PAK-02) were purchased from Cytoskeleton and reconstituted according to the manufacturer’s instructions. Cells were lysed in buffer containing 50 mM Tris-HCl (pH 6.8), 150 mM NaCl, 1% Triton X-100, 20 mM MgCl₂, and 2× protease and phosphatase inhibitors (Thermo Fisher Scientific, 78442). Lysates were clarified by centrifugation at 14,000 rpm for 10 minutes at 4°C, and protein concentrations were normalized using the Pierce BCA Protein Assay Kit (Thermo Fisher Scientific, 23225). For pull-down assays, 50 μg of GST-tagged PAK-PBD was added to each lysate and incubated for 30 minutes at 4°C. Beads were then washed and proteins were eluted in boiling 2× sample buffer (0.2 M Tris-HCl, pH 6.8, 4% SDS, 20% glycerol, 0.03% bromophenol blue, and 5% β-mercaptoethanol).

### Immunoprecipitations

Confluent monolayers of cells were rinsed with cold PBS and then lysed in cold lysis buffer containing 25 mM HEPES (pH 7.5), 200 mM NaCl, 1% Triton X-100, 5 mM MgCl₂, and 2× protease and phosphatase inhibitors (Thermo Fisher Scientific, 78442). Lysates were passed through a 21G needle five times, clarified by centrifugation at 14,000 rpm for 10 minutes at 4°C, and protein concentrations were normalized using the Pierce BCA Protein Assay Kit (Thermo Fisher Scientific, 23225). Equalized lysates (by protein content and volume) were incubated with Dll4 (D7N3H, Cell Singaling Technology, 1:200 IP), HA-tag (C29F4, Cell Singaling Technology, 1:200 IP), S1PR1 (E8U3O, Cell Signaling Technology)1:200 IP) overnight at 4°C with end-over-end rotation. Immune complexes were bound using Protein A beads (Life Technologies) under the same conditions, then washed three times with lysis buffer, and then eluted in 2× sample buffer (0.2 M Tris-HCl, pH 6.8, 4% SDS, 20% glycerol, 0.03% bromophenol blue, and 5% β-mercaptoethanol) and analyzed by SDS-PAGE and immunoblotting using chemiluminescent HRP detection. Western blot images were adjusted for brightness and contrast using ImageJ.

### Notch1 reporter experiments

A Notch reporter line of CHO T-REx1 cells stably expressing 12xCSL-mCitrine, full-length Notch1, and H2B-BFP (a kind gift from Dr. John Ngo, Boston University) was used for all reporter assays. For co-culture experiments, reporter cells were seeded at a 1:1 ratio with either engineered CHO sender cells or wild-type MVECs and cultured for two days. Expression of S1PR1 and Notch ligands was induced with 250 nM doxycycline 24 hours prior to S1P addition.

Time-lapse imaging was performed every 30 minutes for 18 hours using a 10× objective on a Zeiss Axiovert inverted microscope equipped with a Photometrics Prime 95B 22 mm scientific CMOS camera and Zen 2.3 Pro software (Blue Edition, Zeiss). Cell tracking was automated using the TrackMate-StarDist plugin in ImageJ by segmenting BFP-expressing nuclei. The rolling average of nuclear Notch reporter fluorescence (YFP) was calculated over time and normalized to the intensity at the time of S1P addition for each condition. Mann–Whitney U tests were used to compare reporter intensity at the 18-hour timepoint, which showed significant differences between S1P-treated and untreated conditions. This timepoint was therefore used for endpoint analysis in subsequent Notch reporter assays.

As the 18-hour timepoint yielded significant differences in S1P and untreated conditions, we used this timepoint and completed endpoint assays for subsequent experiment. In these endpoint assays, images were acquired using the same Zeiss Axiovert microscope setup. At least ten fields of view (FOVs) per condition were imaged, and mean YFP fluorescence intensity was measured within segmented areas corresponding to BFP-expressing nuclei. Mean reporter intensity per FOV was normalized to the untreated condition. Normalized values across all images are displayed as box-and-whisker plots, with the average from each experiment overlaid as individual data points. Welch’s t-tests were used to compare the mean experimental reporter intensities.

### Statistical Information

Statistical analyses were performed using GraphPad Prism. Comparisons between two groups were analyzed using unpaired, two-tailed Student’s *t* tests assuming unequal variances, unless otherwise indicated. For the time-lapse imaging dataset involving reporter intensities, statistical significance was assessed using the Mann–Whitney *U* test due to its non-normal distribution. Bar graphs representing permeability assays show mean ± standard deviation (SD). All other bar graphs report mean ± standard error of the mean (SEM). Box-and-whisker and violin plots represent the full data range and are overlaid with dot plots where each dot corresponds to an independent experimental average, and the horizontal line indicates the mean of those averages. All images shown are representative of at least three independent experiments

