## Supplemental Figures for "Sphingosine-1-phosphate cross-talks to Notch via a S1PR1-Dll4-MPDZ complex to regulate endothelial barrier function"

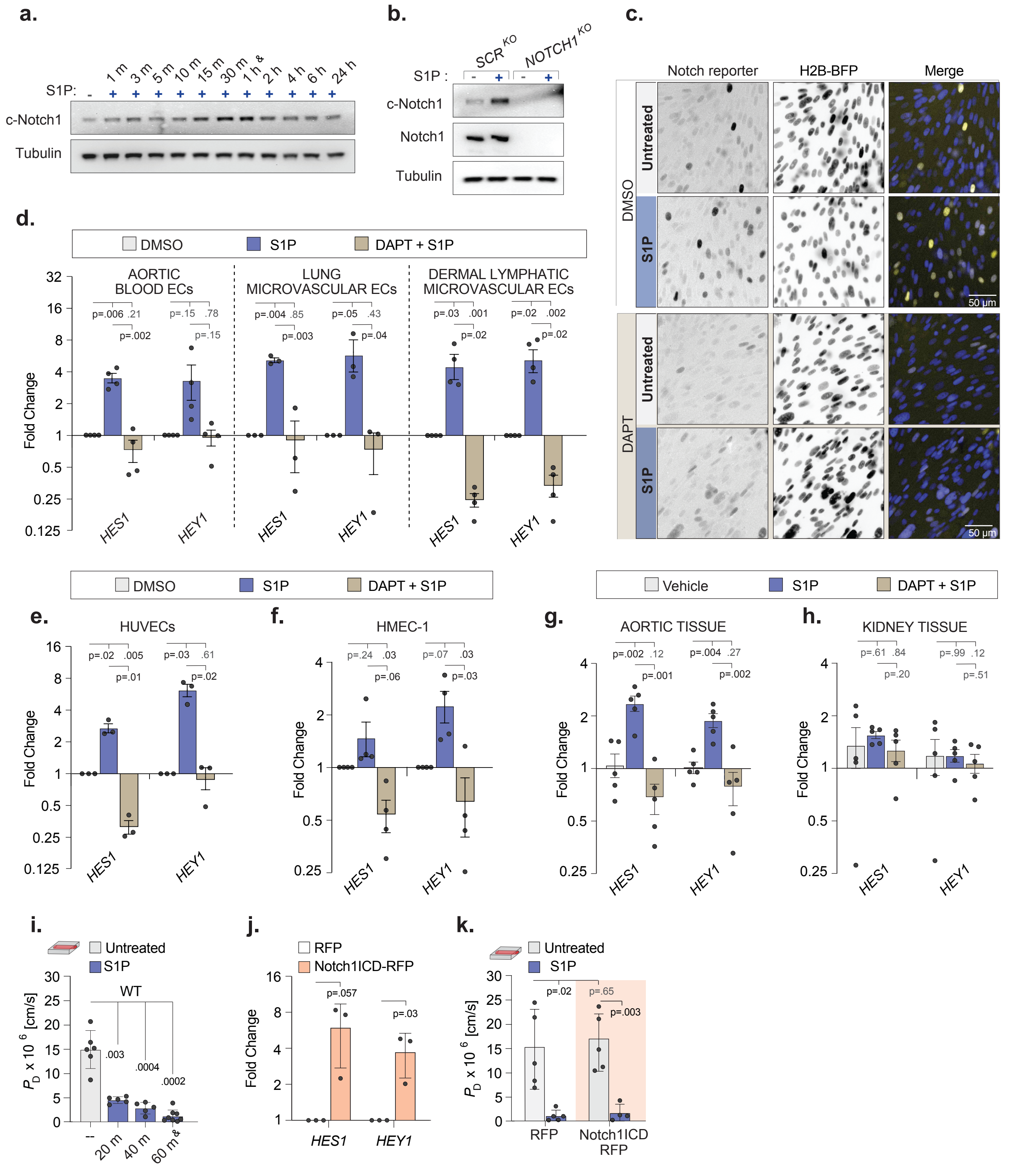


Supplemental Figure 1. S1P induces Notch1 activation in multiple endothelial cells and tissues to regulate vascular barrier independent of Notch transcription.

a, Notch1 cleavage in MVEC lysates following exposure to 100 nM S1P for the indicated time intervals (m, minutes; h, hours). Cleaved Notch1 (c-Notch1) and total Notch1 were assessed by western blot. The selected time point (1 hour) for subsequent assays is marked (&). **b,** Notch1 cleavage in lysates of *SCR KO* versus *NOTCH1 KO* MVECs treated with or without S1P. **c,** Fluorescent micrographs of YFP Notch1 reporter expression (yellow) following S1P exposure, with or without DAPT; nuclei marked with H2B-BFP (blue). Scale bar, 50 µm. **d,** Fold change in *HES1* and *HEY1* gene expression levels in tissue-derived primary endothelial cells, from aortic blood endothelial cells (ECs), lung microvascular ECs and dermal lymphatics (mean ± SEM). **e-f,** Fold change in *HES1* and *HEY1* gene expression levels in (e) HUVECs and (f) the immortalized microvascular endothelial cell line HMEC-1. **g-h,** Fold change in *HES1* and *HEY1* gene expression levels in mice tissues from (g) descending aorta and (h) kidney tissues isolated from mice intravenously injected with S1P or co-treated with DAPT and S1P, normalized to vehicle-treated controls (n = 5, age- and sex-matched littermates). **i**, Diffusive permeability in engineered microvessels lined with wild-type

Supplemental Figure 1. S1P induces Notch1 activation in multiple endothelial cells and tissues to regulate vascular barrier independent of Notch transcription. (Continued.)

(WT) MVECs treated with S1P for various time points or left untreated. The selected time point (1 hour) for subsequent assays is marked (&). **j**, Fold change in *HES1* and *HEY1* gene expression levels in cells expressing fluorescently tagged Notch1 ICD relative to fluorescent RFP control (mean ± SEM). **k**, Diffusive permeability in vessels lined with cells expressing ICD-RFP versus RFP control cells, with or without S1P treatment. RFP control vessels share values with those in Figure 1v.

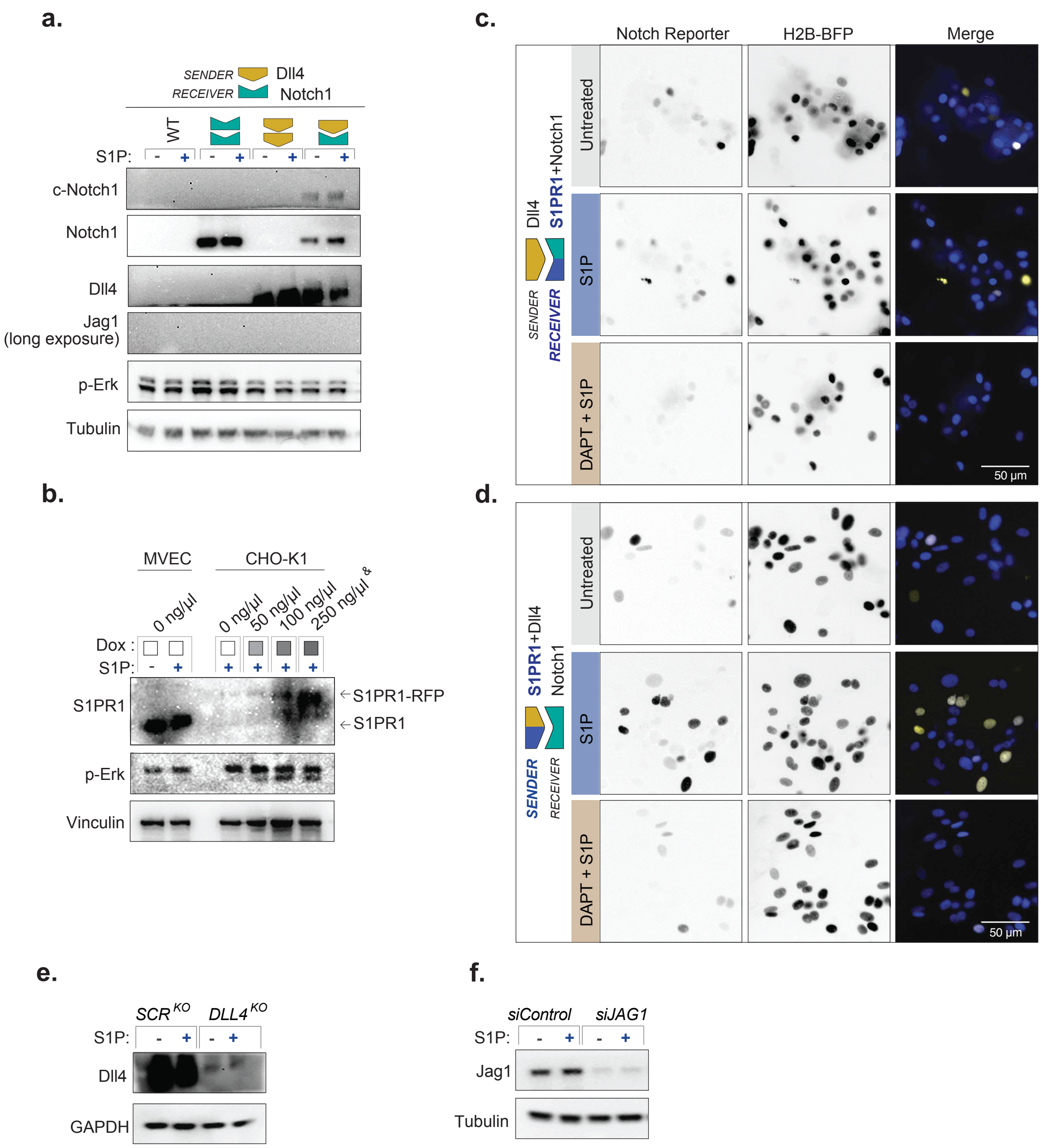


### Supplemental Figure 2. S1PR1 expression enables S1P-induced Notch1 activation in a Dll4-dependent manner.

**a**, Western blot showing expression of Notch1 receptor and ligands and that Notch cleavage and Erk phosphorylation are unchanged by S1P treatment in CHO cells expressing Notch1 or Dll4, either alone or in co-culture. **b,** Immunoblot showing titration of doxycycline to induce expression of S1PR1 in CHO cells to level comparable to endogenous S1PR1 in MVECs. The selected working dose (250 ng/µL) for subsequent assays is marked (&). **c-d,** Micrographs showing Notch reporter intensities in co-cultures of Notch1-expressing receiver cells and Dll4-expressing sender cells with S1PR1 in (c) receivers or (d) senders 18 hours after exposure to S1P, DMSO vehicle, or S1P + DAPT. Scale bar, 50 µm. **e,** Western blot verifying Dll4 knockout (*DLL4 KO*) in MVECs using CRISPR-Cas9, compared to a scrambled control (*SCR KO*). **f,** Immunoblot confirming Jag1 knockdown in MVECs treated with Jag1-targeting siRNA (*siJAG1*) versus a non-targeting control (*siControl*).

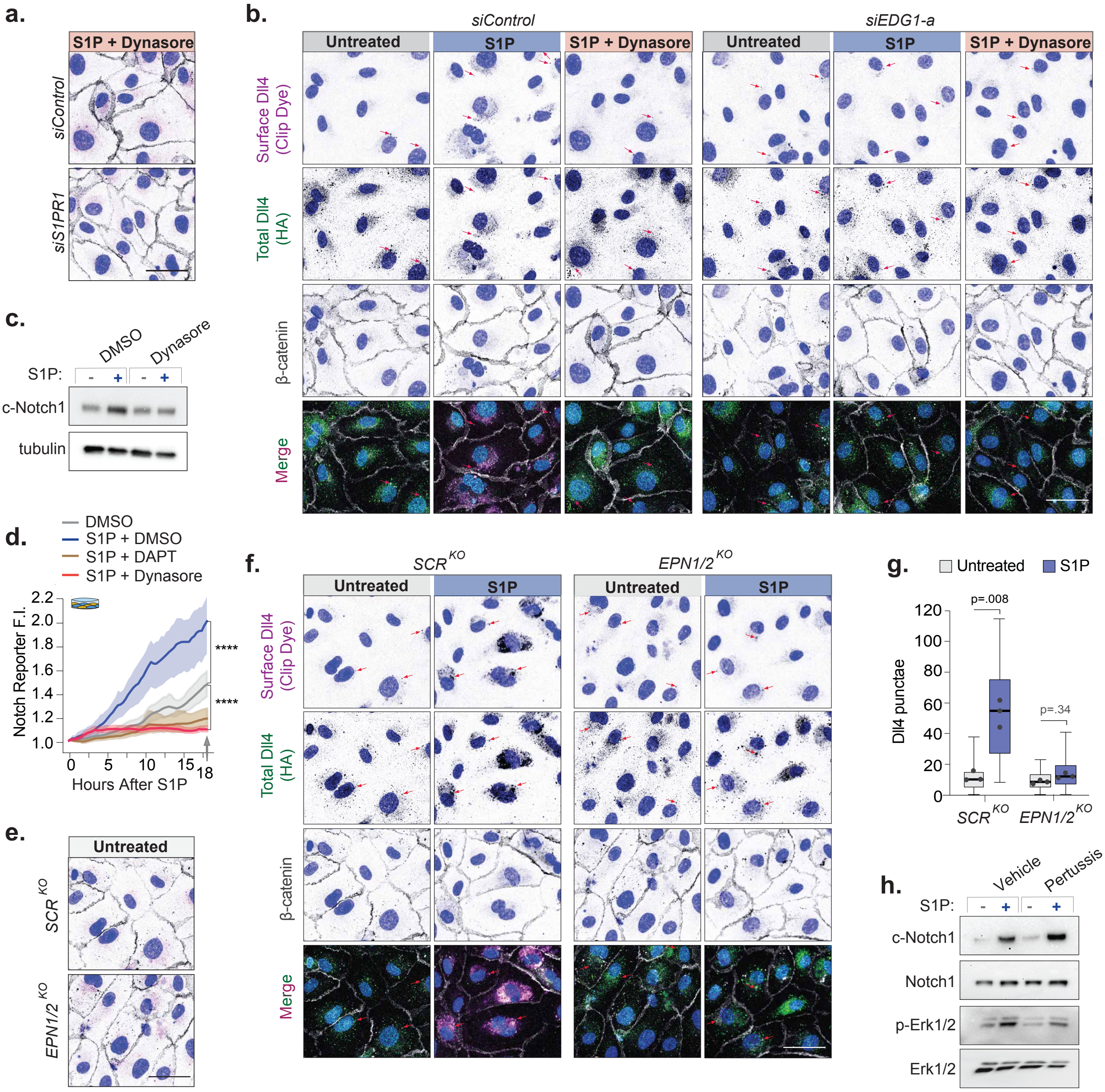


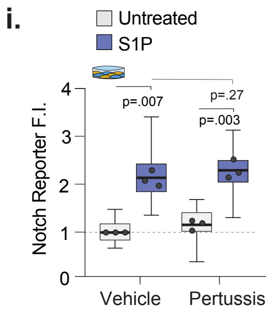


### Supplemental Figure 3. Inhibition of dynamin and epsin impairs Dll4 internalization and suppresses Notch1 activation.

**a,** Merged florescent images of MVECs expressing a dual-tagged (HA and CLIP) Dll4 construct, treated with *siEDG1-a* (S1PR1-targeting siRNA) or *siControl*, followed by treatment with the dynamin inhibitor Dynasore or DMSO vehicle ± S1P, as shown in Figure 3A. Prior to S1P treatment, surface-exposed Dll4 was labeled with a cell-impermeable CLIP dye (magenta). After fixation and permeabilization, cells were stained with anti-β-catenin (gray) to mark cell boundaries and Hoechst (blue) to label nuclei. Scale bars, 50 μm. **b,** Individual fluorescent channels corresponding to the merged images in Figures 3A and S3A. After fixation and permeabilization, cells were stained with anti-HA (green) to visualize total Dll4, anti-β-catenin (gray) to mark cell borders, and Hoechst (blue) to label nuclei. Scale bars, 50 μm. **c,** Immunoblot of MVEC lysates treated with Dynasore or DMSO (vehicle) with or without S1P. **d,** Line graphs showing Notch reporter activation over time in CHO co-cultures treated with DMSO, S1P, S1P + DAPT, or S1P + Dynasore. Statistics done at 18 h mark. **e,** Merged Fluorescent micrographs corresponding to untreated conditions in Figure 3C. Scale bars, 50 μm. **f,** Florescent micrographs of *SCR KO* and *EPN1/2 KO* expressing a dual-tagged (HA and CLIP) Dll4 construct, ± S1P. **g,** Quantification of internalized surface-labelled Dll4 in SCRKO and EPN1/2KO MVECs ± S1P. **h,** Immunoblot of MVEC lysates treated with pertussis toxin or vehicle, ± S1P. **i,** Box and whisker blot of Notch1 reporter fluorescence intensity (F.I.) following pertussis or vehicle treatment, ± S1P. Data normalized to vehicle/untreated controls.

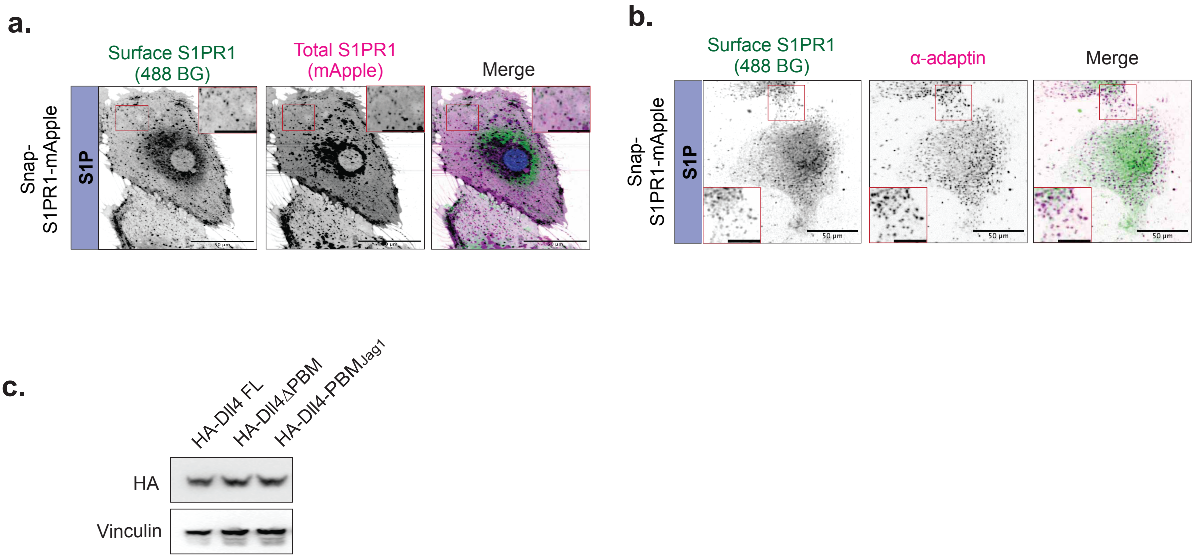


### Supplemental Figure 4. Validation of S1PR1 and Dll4 PBM constructs.

**a,** Fluorescence images of MVECs expressing SNAP-S1PR1-RFP, labeled with a cell-impermeable SNAP dye (green) before S1P treatment to mark surface S1PR1, compared to total S1PR1 (magenta). Scale bars, 20 μm. **b,** Fluorescence images of MVECs expressing SNAP-S1PR1, labeled for surface S1PR1 before S1P treatment, and immunostained for the endocytic protein α-adaptin. Enlarged regions (2× magnification) are shown in red boxes. Scale bar, 20 μm. **c,** Immunoblot showing lysates from CHOs expressing either hemagglutinin (HA)-tagged full-length Dll4 (Dll4 FL), or a modified Dll4 lacking its PDZ binding motif (Dll4ΔPBM) or in which its PBM was replaced with the PBM from Jagged1 (Dll4-PBMJag1).
